# Excessive cortical beta oscillations are associated with slow-wave sleep dysfunction in mild parkinsonism

**DOI:** 10.1101/2023.10.28.564524

**Authors:** Ajay K. Verma, Bharadwaj Nandakumar, Kit Acedillo, Ying Yu, Ethan Marshall, David Schneck, Mark Fiecas, Jing Wang, Colum D. MacKinnon, Michael J. Howell, Jerrold L. Vitek, Luke A. Johnson

## Abstract

Increasing evidence associates slow-wave sleep (SWS) dysfunction with neurodegeneration. Using a within-subject design in the nonhuman primate model of Parkinson’s disease (PD), we found that reduced SWS quantity in mild parkinsonism was accompanied by elevated beta and reduced delta power during SWS in the motor cortex. Our findings support excessive beta oscillations as a mechanism for SWS dysfunction and will inform development of neuromodulation therapies for enhancing SWS in PD.

---

Slow-wave sleep (SWS), the deepest stage of non-rapid eye movement (NREM) sleep, plays a crucial role in learning and memory consolidation^1,2^. Dysfunction of SWS in people with advanced Parkinson’s disease (PD) is associated with faster disease progression^3^, cognitive impairment^4,5^, and daytime fatigue^6^, and is a major factor impacting the quality of life of people with PD. Whether SWS is also disrupted (reduction in SWS quantity) in the early disease state (i.e., mild parkinsonism) and the potential neural mechanism(s) driving dysfunction of SWS in mild parkinsonism remain unclear. An improved understanding of the pathophysiological changes associated with SWS dysfunction in mild parkinsonism can inform the development of targeted therapies to improve SWS in early PD and potentially slow disease progression^7^.

Excessive beta (8-35 Hz) oscillations in the basal ganglia-cortical circuit are considered by many to represent a neural biomarker associated with PD motor signs and are increasingly used to inform closed-loop deep brain stimulation (DBS) approaches^8,9^. Recent evidence suggests that beta oscillations may also play a role in sleep-wake disturbances in advanced parkinsonism^10–14^. Several studies have also shown that cortical delta oscillations, a hallmark of SWS, are reduced in an advanced stage of parkinsonism^4,5^. Together these findings provide compelling support for investigating if the power of cortical delta and beta oscillations is altered in mild parkinsonism and whether they underlie SWS dysfunction.

To elucidate the effect of mild parkinsonism on SWS quantity (% of recording time) and cortical neural oscillations in delta (0.5-3 Hz) and beta (8-35 Hz) bands, we recorded local field potentials (LFPs) from the primary motor cortex (MC) in a nonhuman primate (NHP) across normal and mildly parkinsonian sleep. The MC is a critical node in the basal ganglia thalamocortical network^15^ and exhibits pathological oscillations with depletion in dopamine tone^16–18^. Furthermore, MC displays high amplitude slow oscillations during SWS^19^, a neural characteristic of SWS, hence a relevant recording site to understand the pathophysiology of SWS dysfunction associated with parkinsonism. The goal of this study was to provide insight regarding alterations in cortical neural oscillations as they relate to SWS dysfunction in an early stage of parkinsonism. The findings of this study will be critical for informing the development of targeted neurostimulation therapies to enhance neural oscillations in the cortex that can increase SWS quantity while suppressing neural oscillations detrimental to SWS.

The data reported in this study are from the same nonhuman primate (NHP) in normal and mildly parkinsonian states (see Methods). An example of overnight spectrograms highlighting the reduction in delta oscillations and elevation in beta oscillations during sleep in the mildly parkinsonian state is shown in **Figure 1a** and **Figure 1b**, respectively. The PSD (median±mad) across normal and parkinsonian sleep demonstrates that in the mild parkinsonian state, there was a decrease in delta and increase in beta oscillations in the MC during SWS, **Figure 2a**. Additionally, in mild parkinsonism we observed that SWS quantity was reduced (p=0.027) compared to normal **(Figure 2b)**. The distributions of average delta and beta power during SWS across the recording sessions in normal and parkinsonian states are shown in **Figures 2c** and **2d**, respectively. The power of delta oscillations was reduced (p=0.038) during SWS in the parkinsonian state compared to normal, while the power of beta oscillations was elevated (p=0.001) during SWS. Furthermore, MC delta power positively correlated (r=0.43; p=0.037) with SWS quantity **(Figure 2e)**, while beta power negatively correlated (r=-0.65; p<0.001) with SWS quantity **(Figure 2f)**, suggesting elevated beta oscillations during SWS can be detrimental for SWS. Lastly, mild parkinsonism-associated cortical power changes (delta and beta) during SWS point towards the utility of MC neural oscillations for early-stage disease classification **(Figure 2g)**.

**Figure 1.**
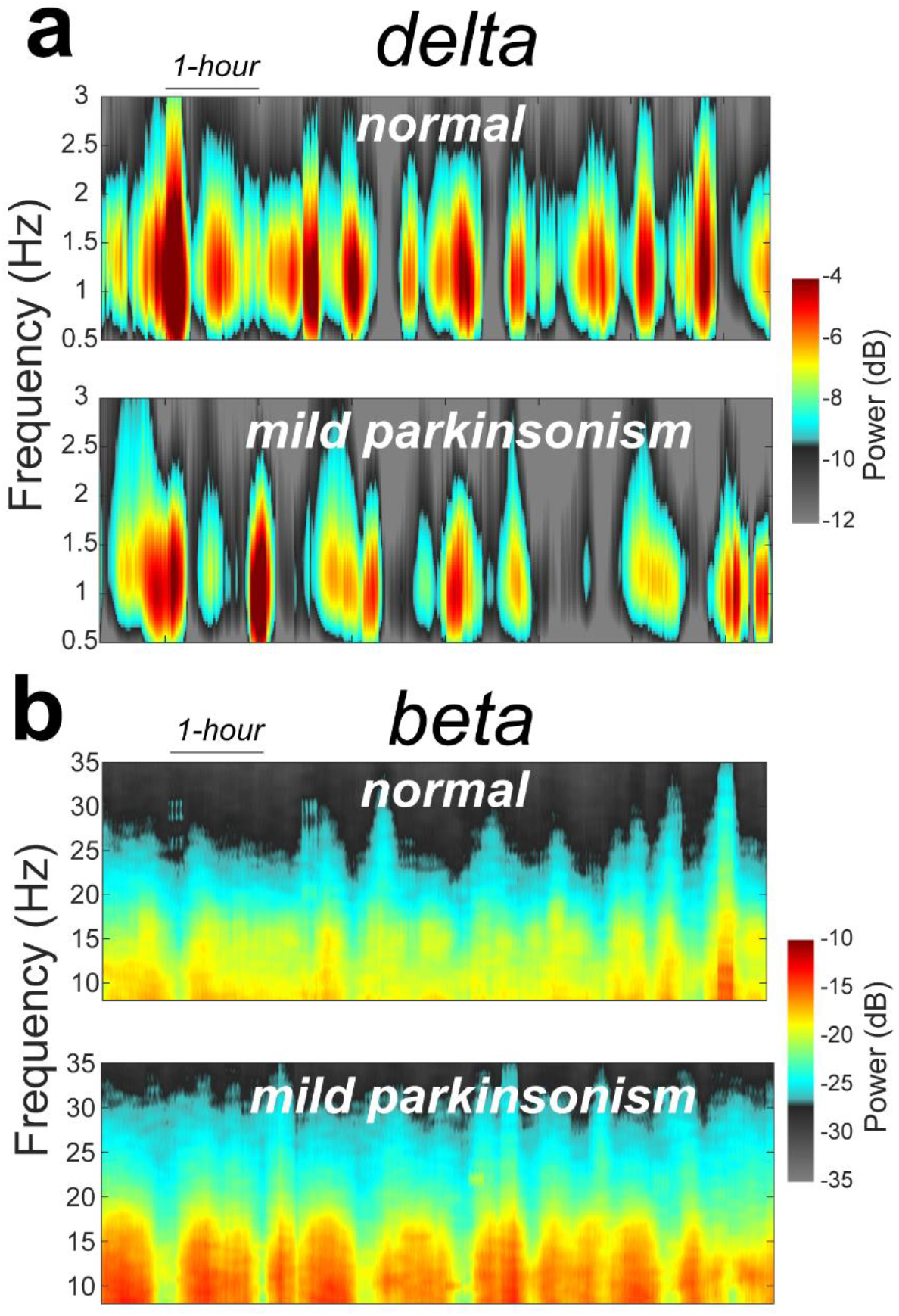
An example of an MC spectrograms for one session of overnight sleep recording in normal and mildly parkinsonian state highlighting parkinsonism-related changes in delta **(a)** and beta **(b)** frequency bands.

**Figure 2.**
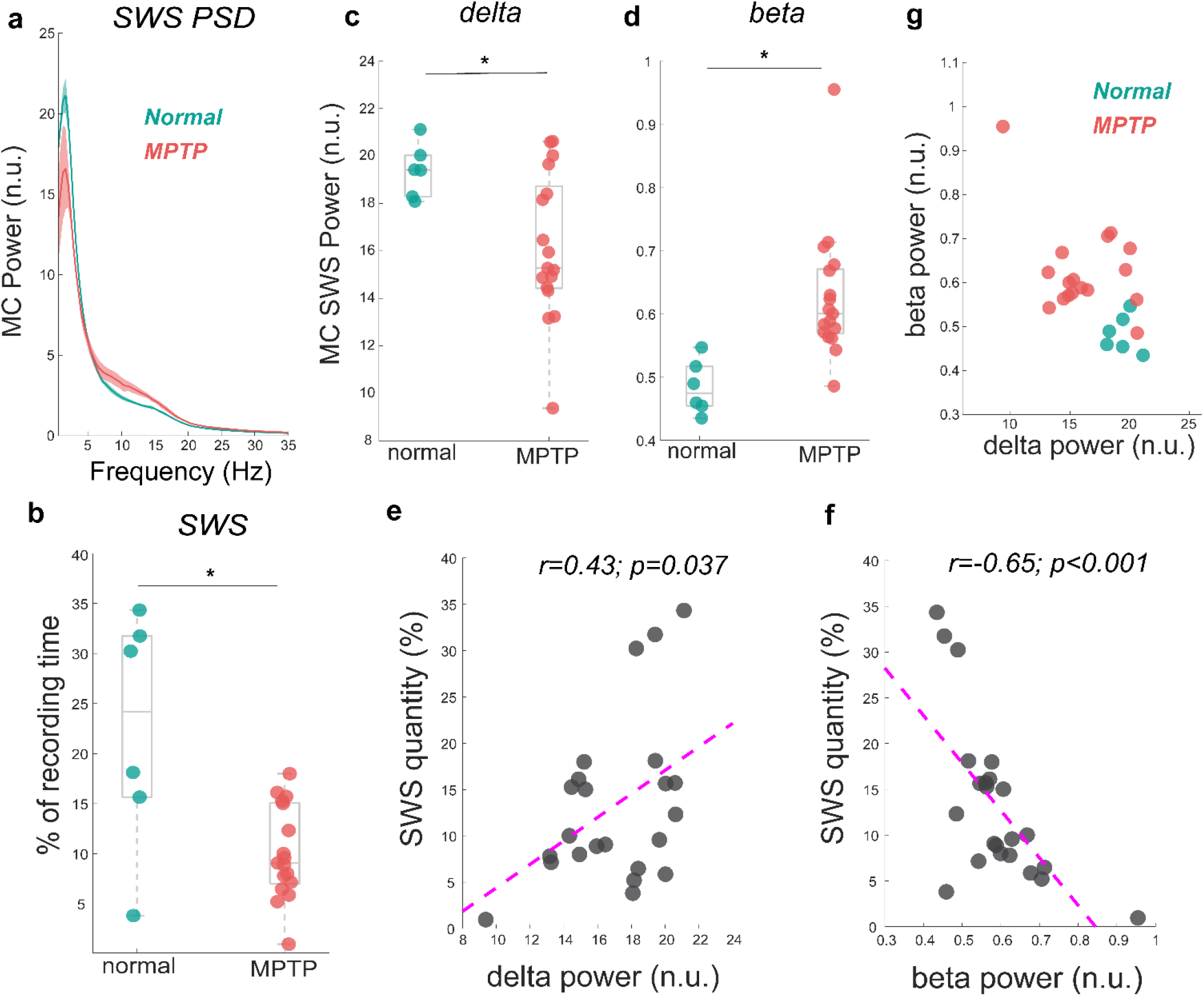
The MC PSD during SWS across normal and mild parkinsonian states showing decrease in delta but an increase in beta power **(a)**. The reduction in SWS quantity (% of the recording time) in mildly parkinsonian state compared to normal is shown in **b**. The distribution of delta and beta power across normal and parkinsonian night showing reduction in delta power (p=0.038) but an elevation in beta power (p=0.001) is shown in **c** and **d**, respectively. The MC delta and beta power correlated significantly with SWS quantity, this correlation is summarized (n=23) in **e** and **f**, respectively. The utility of delta and beta oscillations during SWS for differentiating mild parkinsonism from normal state is characterized in **g**. * Represents significant difference between normal and parkinsonian conditions.

Motivated by previous studies that demonstrated the adverse effects of SWS dysfunction on the quality of life of people with advanced PD^4,5^, we sought to understand the pathophysiology of SWS dysfunction in mild parkinsonism. The key finding of this study was that a marked reduction in SWS quantity in mild parkinsonism was accompanied by excessive beta oscillations and a reduction in delta oscillations in the MC during SWS. Our results suggest that the presence of excessive beta oscillations in the MC during SWS in mild parkinsonism has a detrimental effect on sustaining slow cortical oscillations which may contribute to the reduction in SWS quantity we observed in mild parkinsonian state.

In addition to their recognized role in the development of PD motor signs^20^, our findings provide further support for the role of beta oscillations in the sleep dysfunction observed in patients with PD^11,13^. It has recently been hypothesized that parkinsonism-related excessive beta oscillations in the basal ganglia are transmitted to the cortex, disrupting the initiation or maintenance of slow wave oscillations during sleep^11,21^, though the exact mechanisms by which this occurs remain unclear. Additional investigations utilizing simultaneous recordings at the single neuron and field potential level across the basal ganglia-thalamocortical (BGTC) network will be needed to further test this hypothesis and provide key insight into the neuronal mechanisms driving the disruption of cortical delta oscillations in parkinsonian sleep and the role beta oscillations in the BGTC network may play in the emergence of sleep dysfunction in PD.

While several human and NHP studies have demonstrated that SWS is disrupted in advanced parkinsonism^11,22,23^, whether SWS dysfunction occurs during the early phase of dopaminergic system degeneration is not clear in the literature. A comparison of SWS in idiopathic REM sleep behavior disorder (RBD, often a prodromal stage of PD) patients and healthy controls have shown variable results^24–28^. A study comparing healthy controls, de-novo PD patients, and PD patients with dopaminergic treatment reported a reduction in SWS quantity in the treatment group but not in de-novo PD patients^29^. Another study found no difference in SWS quantity among controls, de-novo PD, and people with advanced PD^30^. Typically, clinical sleep studies are limited to single-night polysomnography in an unfamiliar environment and lack multiple sessions of sleep recordings required to capture the variability in sleep quantity. These limitations could be a contributing factor to the variability reported in the literature as it relates to SWS dysfunction. In this regard, sleep studies using the NHP model of PD can be useful in improving pathophysiological understanding of sleep-wake dysfunction by characterizing within-subject changes with increasing disease severity.

The present study provides support for further investigation into understanding the alterations in SWS in the early stages of parkinsonism. The neural correlates of disrupted SWS (i.e., reduction in cortical delta power and elevation in beta power) that we found may translate to an early screening of PD-related disruption in SWS **(Figure 2g)**. Furthermore, our findings provide a neural basis for the development of targeted therapies for enhancing SWS in people with early-stage PD and potentially slowing disease progression^3,7^. Targeted therapy in the early stage of parkinsonism may include, for example, non-invasive neuromodulation techniques^31,32^ for selective amplification of delta oscillations that can enhance SWS and suppression of beta oscillations that are detrimental to SWS

A major limitation of this study was the limited sample size. Future studies are warranted to reproduce and expand the findings reported here. The ability to perform multiple sessions of sleep recordings in the MC across normal and parkinsonian states is unique to the NHP preclinical studies and provides preliminary insights into the neural correlates of SWS dysfunction in mild parkinsonism. Future preclinical studies that explore neuromodulation (e.g., both non-invasive as well as DBS^33^) techniques for suppressing beta oscillations during sleep and characterizing its effect on cortical SWS to further support the notion that suppression of excessive beta oscillations during sleep can profoundly improve sleep quality. Moreover, simultaneous neural recordings across the basal ganglia thalamocortical network and directed connectivity analysis can help decipher the mechanisms by which beta oscillations in the cortex are elevated during SWS in the early stage of parkinsonism^11,21,34^. Lastly, a higher number of sleep recordings across normal and early PD along with machine learning algorithms will be necessary to generalize the efficacy of features discussed in this study for early-stage disease classification.

## Methods

### Experimental Protocol and Data Collection

All procedures were approved by the University of Minnesota Institutional Animal Care and Use Committee and complied with the US Public Health Service policy on the humane care and use of laboratory animals. One adult female rhesus macaque NHP (23 years old) was used in this study. The subject was instrumented with a 96-channel Microdrive (Gray Matter Research) with microelectrodes targeting basal ganglia, motor thalamus, and motor cortices. A subset of microelectrode channels (n=9) in the MC that were not moved during normal and mild parkinsonian states were used for characterizing the effect of mild parkinsonism on SWS neurophysiology.

The video and wireless local field potential recording from MC during sleep-wake behavior were obtained while the subject was in its home enclosure using a Triangle BioSystem International (TBSI) and Tucker Davis Technology (TDT) recording systems across the normal and parkinsonian state at a sampling rate of ∼24000 Hz. The subject was rendered mildly parkinsonian by administering four weekly low-dose (0.3 mg/Kg) intramuscular injections of the neurotoxin 1-methyl-4-phenyl-1,2,3,6-tetrahydropyridine (MPTP). After the 4^th^ MPTP injection, the subject demonstrated mild motor impairment. After this point, the subject’s motor assessment was routinely monitored for stability for one month, after which the sleep recordings were performed. The severity of the subject’s parkinsonism on the side contralateral to the electrode implants was assessed using the modified version of the Unified Parkinson’s Disease Rating Scale (mUPDRS), which rates symptoms of bradykinesia, akinesia, rigidity, and tremor of the upper and lower limbs as well as food retrieval on a scale of 0-3 (0=normal, 1=mild, 2=moderate, and 3=severe), maximum score = 27 (adapted from Wang et al., 2022^35^). A composite score of 3-9 is considered mild, 10-18 is moderate, 19-27 is severe parkinsonian state. The results presented in the manuscript are from 23 sessions of sleep recordings (6 normal and 17 mildly parkinsonian, mUPDRS=4.61±1.50, mean±SD). The sleep recordings began at approximately 7 pm (lights off at 6 pm) and resulted in 7.66 hours (median value) of recording across normal and parkinsonian states presented in this study.

### Data Analysis

Neural activities recorded from 9 adjacent microelectrode channels were averaged to derive MC LFP. The resultant MC LFP was bandpass filtered from 0.5-700 Hz then resampled to ∼200 Hz and normalized to have unit standard deviation before further processing. Using the Welch power spectral density (PSD) the median power of MC delta oscillations (0.5-3 Hz) was computed on a second-by-second basis with a window size of 128 samples, 50% overlap, and 512 FFTs resulting in a frequency resolution of 0.39 Hz. The epochs were identified as slow wave sleep if the epoch was free from movement and the power of MC delta oscillations was greater than four times the power of MC delta oscillations during the wake. The threshold for wake was determined by analyzing a 1-minute movement-free eyes-open segment (determined using video monitoring of NHP behavior).

The SWS quantity for each night was determined as a percentage of the recording time the subject exhibited SWS. The PSD associated with each SWS epoch was normalized by dividing the PSD by total power. The total power underlying the PSD of the respective epoch was obtained by integrating the PSD from 0.5-100 Hz. From the normalized PSD, the median power underlying delta (0.5-3 Hz) and beta (8-35 Hz) bands for each SWS epoch were determined. The average of SWS delta and beta power for each night was obtained for characterizing the effect of parkinsonism on cortical neural oscillations and for correlating with SWS quantity.

### Statistical Analysis

Normality was not assumed, and the Wilcoxon rank-sum (WRS) test was performed to report statistical differences. Reported p-values are an outcome of the WRS test unless stated otherwise. The Pearson correlation coefficient was used to assess the correlation between delta or beta power and SWS quantity. Statistical tests were performed using the statistical toolbox of MATLAB (Mathworks Inc., Natick, MA). The test results were considered significant at p<0.05.

## Authors Contribution

AKV and LAJ conceived the research. AKV, BN, and LAJ developed an analysis methodology. YY, EM, and LAJ performed data acquisition. AKV performed data analysis. AKV, KA, BN, YY, DS, MF, JW, CDM, MJH, JLV, and LAJ interpreted the results. AKV wrote the manuscript and created figures. All authors critically edited the manuscript and approved it for submission.

## Data Availability

Data supporting the findings of this study are available from the corresponding author upon reasonable request.

## Acknowledgment

This work was supported by the National Institutes of Health, National Institute of Neurological Disorders and Stroke (NINDS) R01-NS110613, R01-NS131371, R01-NS058945, R37-NS077657, P50-NS123109, P50-NS098573, MnDRIVE (Minnesota’s Discovery Research and Innovation Economy) Brain Conditions Program, MNREACH, and the Engdahl Family Foundation.

